# The trajectory of cortical GABA levels across the lifespan: An individual participant data meta-analysis of edited MRS studies

**DOI:** 10.1101/2020.07.23.218792

**Authors:** Eric C. Porges, Greg Jensen, Brent Foster, Richard A. E. Edden, Nicolaas A. J. Puts

## Abstract

GABA is the principal inhibitory neurotransmitter in the human brain and can be measured with Magnetic Resonance Spectroscopy (MRS). Conflicting accounts report decreases and increases in cortical GABA levels across the lifespan. This incompatibility may be an artifact of the size and age-range of the samples utilized in these studies. No single study to date has included the entire lifespan. In this study, 8 suitable datasets were integrated to generate a model of the trajectory of GABA across the lifespan. Data were fit using both a log-normal curve and a nonparametric spline as regression models using a multi-level Bayesian model utilizing the Stan language. Integrated data show the lifespan trajectory of GABA involves an early period of rapid increase, followed by a period of stability during early adulthood, with a gradual decrease during adulthood and aging that is described well by both spline and log-normal models. The information gained will provide a general framework to inform expectations of future studies based on the age of the population being studied.

---

Magnetic resonance spectroscopy (MRS) is a non-invasive imaging technique that allows for the measurement of levels of metabolites. Of particular interest to the neurosciences is the measurement of specific neurotransmitters such *γ*-Aminobutyric acid (GABA) in vivo (Alger, 2010; Edden and Barker, 2007; Mescher et al., 1998; Mullins et al., 2014; Puts et al., 2011; Rothman et al., 1993). GABA is the main inhibitory neurotransmitter in the human nervous system and plays a fundamental role in central nervous system function (Buzsáki et al., 2007). A number of studies have explored the relationship between cortical GABA (as measured with MRS) and age in various contexts. These studies have found that age-related changes in GABA are consistently associated with cognitive and neurophysiological outcomes that change across the lifespan. These findings have important implications for both healthy and pathological development and aging.

Most prior studies have included a restricted age range. Some have reported increases in GABA as age increases (Ghisleni et al., 2015), others have reported decreases (Gao et al., 2013; Marenco et al., 2018; Porges et al., 2017b; Rowland et al., 2016; Simmonite et al., 2019), and still others reported no significant age-related changes (Aufhaus et al., 2013; Mikkelsen et al., 2018). This inconsistency makes the results difficult to interpret, as partially overlapping age ranges produce conflicting trajectories. For example, the age range of participants reported by Gao et al. (2013) have substantial overlap with those reported by Ghisleni et al. (2015) yet have disparate outcomes. Interestingly, this apparent conflict in the literature is not present when similar age ranges are considered. For example, both Porges et al. (2017b) and Gao et al. (2013) focus on adults through advanced age and both report age-related decrease in GABA. Here, we predicted that this apparent conflict is the result of a non-linear age-related trajectory, similar to other features of brain development (Lebel et al., 2012), showing that the lifespan trajectory involves a rapid increase of GABA rich grey matter in early life, relative stability in early adulthood, followed by a gradual decrease, and that this is consistent with inhibition-dependent behavior (Williams et al., 1999). To date, no single study has explored the lifespan trajectory of cortical GABA spanning development, adulthood, and aging. In the absence of a lifespan study, we implemented an individual participant data (IPD) meta-analytic approach following PRISMA guidelines (Moher et al., 2009,; see Figure 1) supplemented with data collected by the authors and previously published in summary form (Puts et al., 2017).

**Figure 1.**
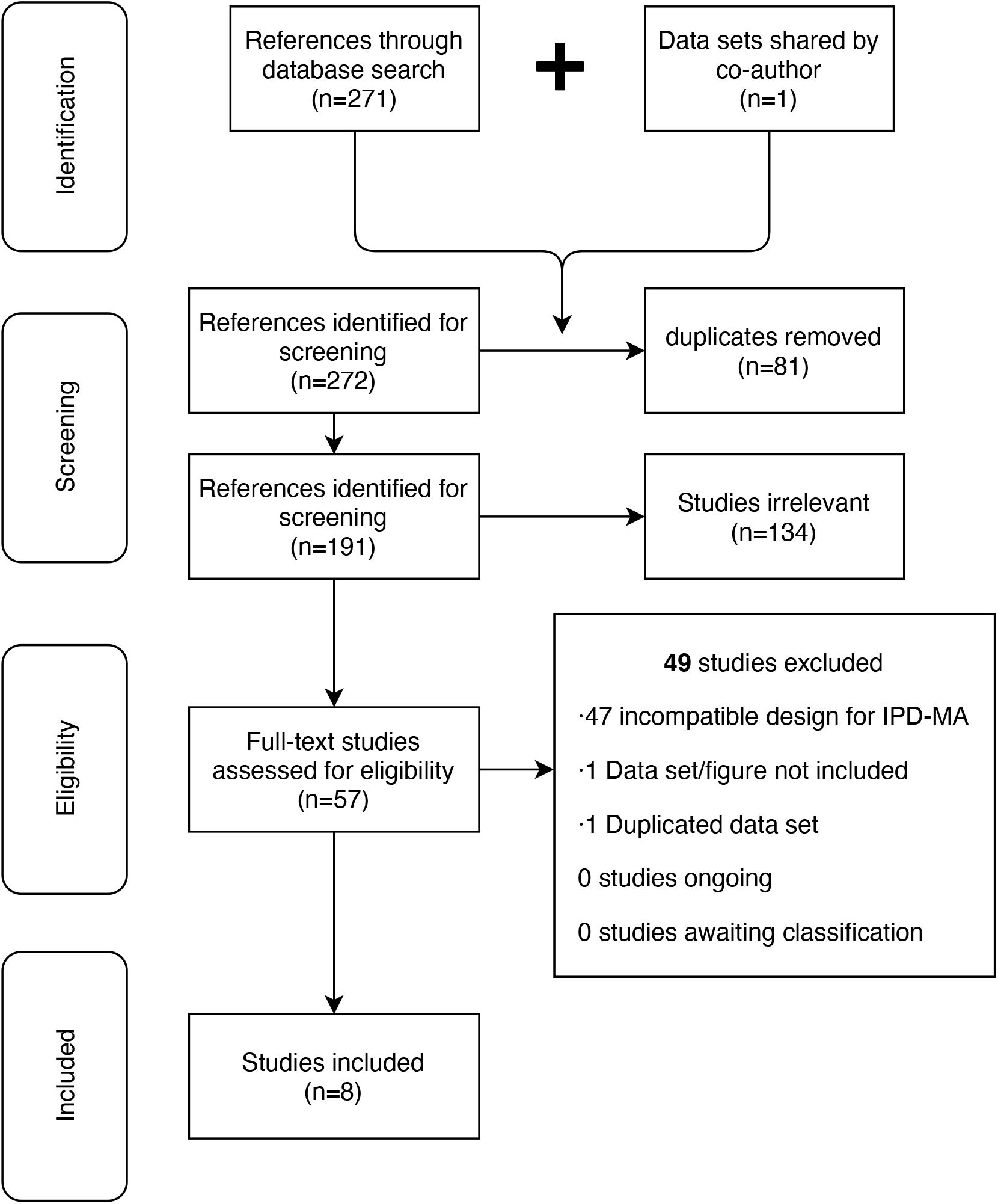
Prisma 2009 Flow Diagram of study identification and inclusion.

The majority of MRS studies of GABA at 3 Tesla have utilized the unique structure of the GABA molecule to selectively edit the GABA signal using a typical MRS acquisition with frequency-selective pulses (e.g. Mullins et al., 2014). Editing is necessary at 3 Tesla due to the low concentration of GABA in the human brain (1-2 mM) (Harris et al., 2017). In un-edited MRS, signal from higher concentration molecules like NAA and creatine (Cr) mask the GABA signal. The most widely used edited MRS technique is MEGA-PRESS (Mescher et al., 1998), in which a GABA-selective editing pulse at 1.9 ppm is applied in half of the experiment (edit-ON). Due to its low concentration, measurement of the GABA-edited signal in humans requires a large voxel (most commonly 27 cm3 Mullins et al., 2014; Peek et al., 2020; Salthouse, 2010) to keep acquisition times reasonable (1̃0 minutes) and to provide an adequate signal-to-noise ratio (Mikkelsen et al., 2018). This limitation constrains the spatial specificity of the measurement to coarse regions that often lack discrete functional specificity. The spectrum is further complicated by a macromolecule signal at 3 ppm, which is coupled to another macromolecule signal at 1.7 ppm and thus falls within the envelope of the editing pulse, resulting in a co-edited macromolecule signal as part of the 3 ppm GABA signal (Edden et al., 2014; Henry et al., 2001). Consequently, most studies refer to the GABA signal as “GABA+”. Both MM-suppressed and GABA+ measures are included here (Table 1).

**Table 1.** Neuroimaging acquisition and analysis details for eight studies included in the analysis. GABA+ = GABA and macromolecule; GABA = Macromolecule-suppressed GABA; H2O = water referenced; Cr = creatine referenced. MRS averages refer to the number of ON+OFF transients. *The manuscript refers to 96 averages. It was clarified with the authors that this referred to 96 ON and 96 OFF averages.

| GABA Study | Type of Scanner | Analysis Software | Reference Method | Voxel Volume (mL) | MRS Averages | TE (ms) | TR (ms) |
| --- | --- | --- | --- | --- | --- | --- | --- |
| Aufhaus* | 3T Siemens | jMRUI / LCModel | GABA/H<sub>2</sub>O | 24 | 192* | 68 | 3000 |
| Gao | 3T Philips | jMRUI | GABA+/Cr | 27 | 320 | 68 | 2000 |
| Ghisleni | 3T GT | LCModel | GABA+/H<sub>2</sub>O | 30 | 320 | 68 | 2000 |
| Mikkelsen | 3T GE / Philips / Siemens | Gannet | GABA+/Cr | 27 | 320 | 68 | 2000 |
| Porges | 3T Philips | Gannet | GABA+/H<sub>2</sub>O | 27 | 320 | 68 | 2000 |
| Puts | 3T Philips | Gannet | GABA+/H<sub>2</sub>O | 27 | 320 | 68 | 2000 |
| Rowland | 3T Philips | Gannet | GABA/H<sub>2</sub>O | 24 | 256 | 68 | 2000 |
| Simmonite | 3T Philips | Gannet | GABA+/Cr | 22.5 | 256 | 68 | 1800 |

The functional relevance of MRS measures of GABA is important to note. Measured GABA levels include both intracellular and extracellular contributions to the overall GABA concentration, however these relative contributions are not well-known, particularly in development. GABA levels measured via MRS at rest describe a physiological characteristic of the tissue measured and, while associated with functional metrics (e.g. neurophysiological response or behavior), are better interpreted in a manner similar to structural neuroimaging. The GABA measure is a single quantity representing the entirety of the voxel and thus, all pools of GABA present (with no discrimination between intra/extracellular, neuronal/astrocytic, vesicular, etc). However, studies have shown that the majority of the GABA measured using MRS reflects intracellular levels rather than synaptic levels (Marenco et al., 2011; Rae, 2014; Stagg et al., 2011b). As such, the GABA signal is considered to be reflective of inhibitory tone rather than dynamic synaptic inhibition (Rae, 2014). GABA levels measured via MRS in young adults at rest have been reported to be stable for up to 7 months (Near et al., 2014) and do not exhibit a diurnal rhythm (Evans et al., 2010).

In this manuscript, we statistically combine datasets of published research that used MEGA-PRESS to measure GABA levels of discrete age ranges where individual data points were presented relative to age to present a non-linear model for GABA levels over the human lifespan using an Individual Participant Data approach. This work is motivated by several studies showing that MEGA-PRESS measures of cortical GABA are relevant to both development and aging with a specific emphasis on cognition and perception. We further discuss the lack of available data across the lifespan.

## The importance of GABA in cognition motivates an understanding of the age-relationship

GABA levels measured with MRS have been linked to clinical and cognitive outcomes. While this review does not pertain to cognition, an understanding of the relationship between GABA and cognition motivates our initial step towards understanding the relationship between GABA and age. Alterations of GABA levels are seen in neurodevelopmental disorders such as ADHD (Bollmann et al., 2015; Edden et al., 2012), Autism Spectrum Disorder (Cochran et al., 2015; Drenthen et al., 2016; Gaetz et al., 2014; Puts et al., 2017), and Tourette syndrome (Puts et al., 2015), as well as in other neurological and psychiatric disorders including schizophrenia (Rowland et al., 2016; Shaw et al., 2020), depression (Sanacora et al., 1999) and neurofibromatosis (Violante et al., 2013). These associations are reviewed by Puts and Edden (2012) and by Schür et al. (2016). Measures of GABA levels appear to be functionally and regionally specific, and studies have shown associations between sensorimotor GABA levels and tactile sensitivity (Heba et al., 2016; Puts et al., 2011, 2015), between occipital GABA levels and visual orientation discrimination (Edden et al., 2009; Yoon et al., 2010), between motor cortex GABA and motor control and learning (Bachtiar and Stagg, 2014; Stagg et al., 2009, 2011a), between anterior cingulate cortex and response inhibition (Silveri et al., 2013), and between frontal GABA concentrations and working memory (Michels et al., 2012). For a review, see Duncan et al. (2014). Moreover, GABA levels correlate with other measures of brain activity, including functional magnetic resonance imaging (fMRI) (Donahue et al., 2010; Muthukumaraswamy et al., 2009), measures of cerebral blood flow (Donahue et al., 2010), and motor cortex gamma oscillations (Gaetz et al., 2011; Muthukumaraswamy et al., 2009) as measured using magnetoencephalography (MEG); for a review, see Duncan et al. (2014).

The above paragraph reflects a large number of studies showing that GABA plays an important role in regulating cognitive function in health and disease. It is also well known that age affects cognitive processes both in development and aging, including sensory processing (Koerner and Zhang, 2018; Simmonite et al., 2019), working memory (Mok et al., 2019), motor function (Hermans et al., 2018; Maes et al., 2018; Mikkelsen et al., 2018) and many others, however the potential role of GABA has not been well-explored. Perhaps more immediately, it is not well-known how GABA changes across the lifespan. Our approach allows for a systematic review of existing work studying GABA across development and aging.

## GABA across the lifespan

To date, cross-sectional and longitudinal investigations of cortical GABA across the entire human lifespan have yet to be published. However, there have been recent reports investigating the relationship between cortical GABA levels and discrete age ranges in humans that test a linear association focused on a specific population (e.g. ‘ aging’). As discussed above, the results of these studies are ambiguous—some suggest a positive correlation between GABA and age, others hint at a negative correlation, and still others conclude that there is no relationship at all. Here, we discuss these reports in the context of human development and divide them into three categories: developmental, adult, and aging.

### Developmental

The developmental component of the lifespan of cortical GABA as measured by MRS of GABA is explored in less depth than in adult or aging cohorts. Port et al. (2017) report a maturational increase in GABA+ with age. The majority of evidence comes from non-MRS work, showing dramatic change in GABAergic function during early life. Human autopsy data describes large changes in both GABA synthesis and receptor expression (Pinto et al., 2010) and animal models describe a shift in GABA from excitatory to inhibitory (Leonzino et al., 2016). Yet, in vivo reports in healthy younger populations (particularly infants and young children) are sparse or missing. Several reasons exist for the absence of high quality MRS data during development with technical challenges being one main challenge. These challenges are not unique to MRS, but exist for most —if not all— Magnetic Resonance modalities. For example, it is well established that imaging of MRS of GABA is highly sensitive to motion (Edden et al., 2016; Mullins et al., 2014), thus compounding the challenges involved when imaging pediatric cohorts. Many studies in pediatric cohorts, including our own (Puts et al., 2017), also suffer from the limitation that age relationships are not reported due to individual studies often studying a restricted age range to minimize developmental effects within the cohort.

### Adult

The vast majority of studies in healthy populations focus on age ranges between development and aging to minimize the effect of age on the measures of interest (and for ease of recruitment). However, this limits the reporting of GABA-age relationships within this range. Mikkelsen et al. (2017) conducted a multisite study collecting GABA+ levels in 272 participants between 18 and 35 years old, providing a substantial dataset to assess this relationship. The sample size and restriction to healthy adults in this study provide a reasonable representation of normal GABA levels in the target demographic. Their objective with this study was to report stability of the GABA+ measure across multiple 3T MRI platforms with systems by GE, Phillips, and Siemens well represented. Voxel placement was selected for the medial parietal lobe. While their original manuscript contains neither a report nor a visualization of the GABA/age relationship, we are able to provide this information for our review (data are freely available from the Big GABA repository, Mikkelsen et al., 2017, https://www.nitrc.org/projects/biggaba/). There was no age-related increase or decrease between age and GABA+/Cr [?2(7) = 3.52, pboot = 0.31] in this large cohort of adults between 18-48 years of age.

### Aging

Most, if not all, MRS of GABA studies that investigate aging populations report a decrease in cortical GABA as a function of age in both frontal (Gao et al., 2013; Porges et al., 2017b) and parietal (Gao et al., 2013) voxels. Marenco et al. (2018) also show a decrease in GABA with aging. It is important to note that other reports (Hermans et al., 2018; Maes et al., 2018) have compared MRS of GABA between defined groups of older and younger adults rather than with continuity across the lifespan. These findings are consistent with continuous approaches, with older adults having reduced GABA. However, a categorical approach comparing two groups does little to elucidate the aging-related trajectory. Manuscripts that employ MEGA-PRESS methodology in a manner that is inconsistent with methods outlined in consensus papers (Mullins et al., 2014; Puts and Edden, 2012), have insufficient SNR, or other technical limitations have not been considered for this assessment.

In conclusion, an understanding of the link between GABA and age is incredibly important for the study of inhibition across the lifespan, the study of development- and aging-related behavioral and cognitive processes, and the study of health and disease. No study has attempted to study GABA across the entire lifespan. Here we utilize an Individual Participant Data Meta-Analytical approach of all existing and eligible cortical edited MRS of GABA data across the lifespan (from development to aging) to build a model that informs us of the best-fit model of GABA across the lifespan. We hypothesize that the model of best-fit would be consistent with that of other cortical measures, showing a sharp increase during development, and a slow gradual decrease during aging.

## METHODS

We conducted and reported this systematic review in accordance with the PRISMA (Preferred Reporting Items for Systematic Reviews and Meta-Analyses) statement (Moher et al., 2009) and we used an Individual Participant Data-Meta Analysis (IPD-MA) approach (Debray et al., 2015).

A systematic literature search was performed in two iterations by BF and EP to retrieve studies in which MRS of GABA using the MEGA-PRESS method was collected in the human brain from voxels that included the frontal lobe. In the first iteration, a search was performed using Google Scholar and Medline with the following combination of terms: (GABA OR GABA+ OR ?-aminobutyric acid OR gamma-Aminobutyric acid) AND (MRS OR Magnetic Resonance Spectroscopy) AND (MEGAPRESS OR MEGA-PRESS OR MEshcher-GArwood Point RESolved Spectroscopy OR edited). Both GABA+ and MM-suppressed measures were deemed inclusive. The following constraints were applied to limit results: the result should be (i) a full-text article or a conference abstract, (ii) peer-reviewed, (iii) written in English, (iv) included in the publication must be a scatter plot with GABA by Age suitable format for extraction of individual human subject data via WebPlotDigitizer. The search was conducted on April 2, 2019, resulting in a total of 273 studies. Out of these, 55 were relevant.

### Additional Criteria

#### Study Design

A second step was performed by BF, EP, and NP to exclude based on the following criteria: studies must report GABA, acquired using MEGA-PRESS, in at least one cortical voxel (subcortical voxels were excluded). To exclude a potential region effect, when a frontal voxel (any inclusion of frontal cortex) was available we included that voxel (given its importance for cognition and applicability to the majority of available studies). If not, we used the other cortical voxel only. If a study sampled multiple voxels, we only extracted data from a frontal voxel to prevent multiple sampling of a single subject as this would include in inclusion of non-independent datasets. If multiple frontal voxels were available, we chose the dataset with the largest sample. Then, data were only included when they originated from published figures of sufficient quality for data extraction of individual data points to be u sed. Finally, the data extracted required contiguous age ranges of 5 or more years for inclusion in the analysis. Studies failing to satisfy all criteria were deemed incompatible with the individual participant data meta-analytic approach (as described below). Duplicate datasets were excluded as well.

#### Data Quality

A third step was performed to assess whether studies adhered to consensus quality assurance criteria for data collection, analysis, and reporting. For this purpose, two coauthors (NP and EP) evaluated all remaining studies using the MRS-Q, which was specifically designed for MRS and based upon consensus documentation (Peek et al., 2020).

Of the 55 relevant studies retrieved in the systematic search, only 7 included figures suitable for data extraction or had data freely available in available online repositories. One dataset was only partly published (Puts et al., 2017) but was supplied in full by authors of this manuscript NP and RAEE, for a total of 8 datasets. At final review, these datasets were reviewed for consistency in research methods (see Figure 1 and Table 1), evaluated for age distributions (see Figure 2 and Table 2), and combined in aggregate (see Figure 3 and Table 3).

**Table 2.** Descriptive statistics for eight studies included in the analysis.

| GABA Study | # of Subjects | Mean Age | Age (SD) | Age Range |
| --- | --- | --- | --- | --- |
| Aufhaus* | 44 | 35.5 | 10 | 21-53 |
| Gao | 96 | 45.7 | 14.5 | 20-76 |
| Ghisleni | 55 | 27.2 | 11 | 13-53 |
| Mikkelsen | 220 | 26.5 | 4.9 | 18-48 |
| Porges | 86 | 71.8 | 10.6 | 43-92 |
| Puts | 101 | 10.3 | 1.2 | 8-13 |
| Rowland | 82 | 38.0 | 13.7 | 18-62 |
| Simmonite | 38 | 50.1 | 29.2 | 18-87 |

**Table 3.** Descriptive statistics for eight studies included in the analysis.

| GABA Study | Mean Slope | Lower Bound (0.025 Quantile) | Upper Bound (0.975 Quantile) | $R^2$ |
| --- | --- | --- | --- | --- |
| Aufhaus* | -0.0008 | -0.0055 | 0.0041 | 0.003 |
| Gao | -0.0075 | -0.0092 | -0.0058 | 0.459 |
| Ghisleni | 0.0034 | 0.0007 | 0.0061 | 0.106 |
| Mikkelsen | -0.0019 | -0.0054 | 0.0016 | 0.005 |
| Porges | -0.0099 | -0.0133 | -0.0064 | 0.279 |
| Puts | 0.0436 | 0.0085 | 0.0795 | 0.058 |
| Rowland | -0.0032 | -0.0060 | -0.0004 | 0.061 |
| Simmonite | -0.0023 | -0.0040 | -0.0005 | 0.156 |

**Figure 2.**
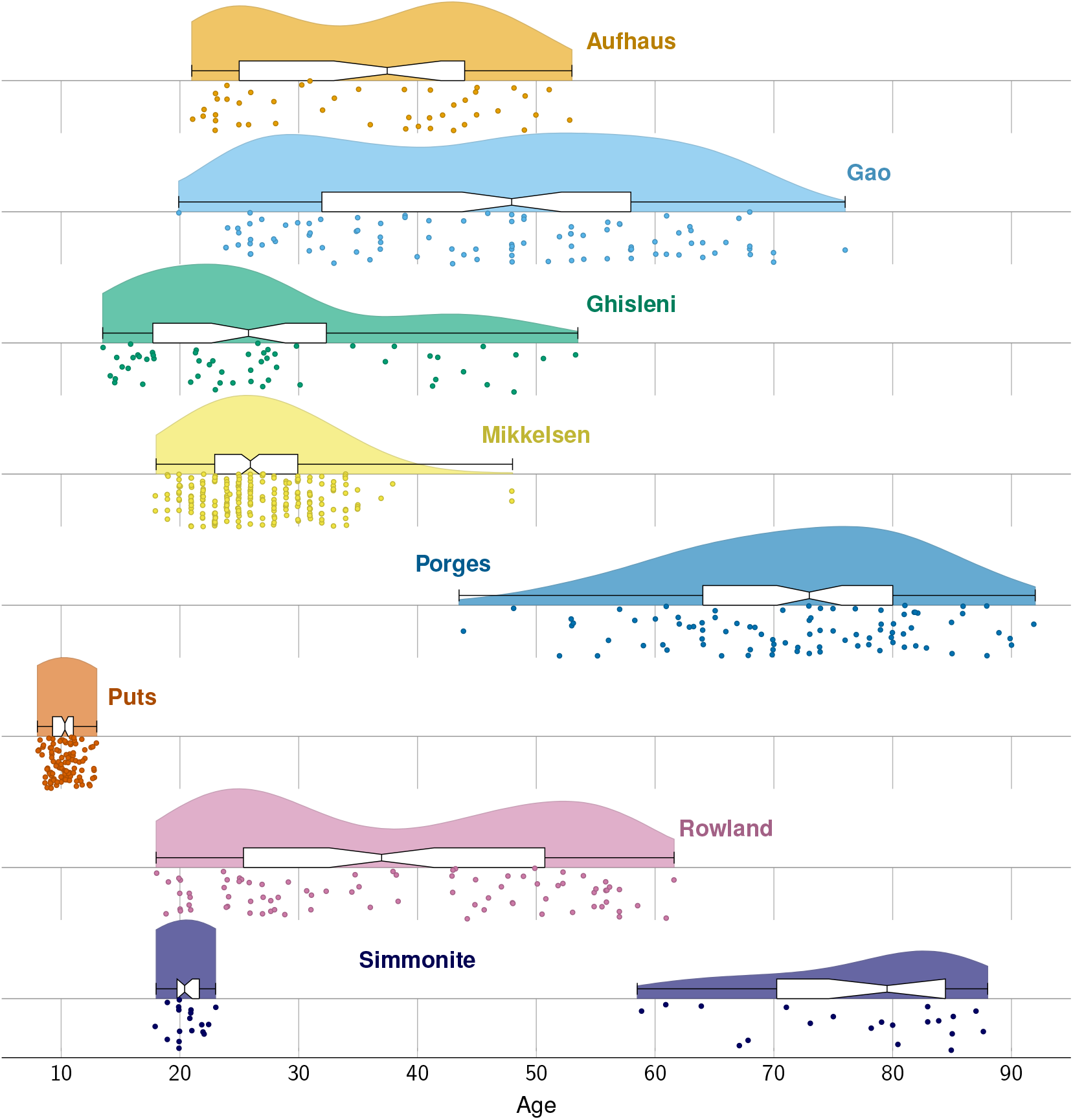
Raincloud plot depicting the age distribution in each of the eight included datasets. Plotted densities are scaled within study. The relevant analysis script for these densities is included as supplemental material.

**Figure 3.**
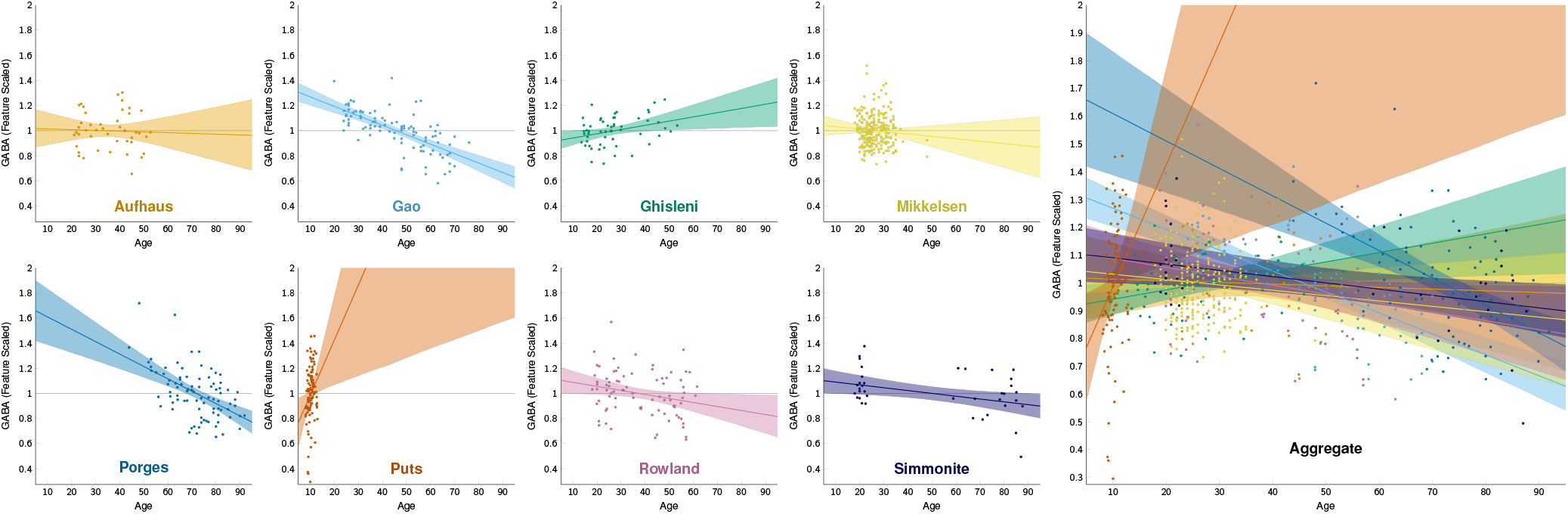
Linear relationships between age and GABA signal. In each dataset, GABA was scaled relative to the geometric mean. Linear models were fit for each dataset separately. Shaded regions represent the 95% credible interval for the regression line.

#### Risk of Bias

All 8 studies included in the systematic review included both male and female participants, although the pediatric data were somewhat skewed to include more males. No data were excluded based on acquisition parameters as assessed with the MRS-Q. Most studies were well-matched for acquisition and followed published recommendations (Mullins et al., 2014). No exclusions were made based on the direction of the correlations. With the exception of two publications, (Gao et al., 2013; Porges et al., 2017b), the age-by-GABA relationship was not a primary outcome, and was therefore unlikely to have been a driver of publication bias in the majority of studies included. However, a risk of publication bias cannot be ruled out.

Individual data points were extracted from figures using WebPlotDigitizer (Rohatgi, 2019). None of the data included were corrected for voxel tissue fractions (see Discussion). MEGA-PRESS sequences can vary between and within MRI vendors; these can impact editing efficiency and in turn absolute quantification (Harris et al., 2015; Saleh et al., 2020). However, the consequence of this will not impact the within-site relationship to age (the metric used in this review) as the consequences of such variation are stable within-site and function as a scaling factor (Mikkelsen et al., 2018).

Table 2 gives a basic description of sample size and age range for the eight datasets. Additionally, Figure 2 depicts the distribution of ages using a raincloud plot (Allen et al., 2019).

### Statistical Methodology

Our meta-analysis made use of an individual participant’s data meta-analytic (IPD-MA) approach (Debray et al., 2015). The chief advantage of this approach is that it allows the analyst to account for the evidence provided by the individual observations recorded in each study while also accounting for any systematic differences between studies using the framework of a hierarchical model. As a result, methodological steps like estimating the “weight” associated with each study in order to determine how to combine reported statistics is rendered unnecessary, as the relative effect size and uncertainty of each study is communicated to the model by the data themselves (Riley et al., 2010). Such an approach is especially important in non-linear regression paradigms, in which the meta-analytic models’ uncertainty for any value of a continuous predictor depends on the complex covariance of participant-level and study-level parameters. Without considering the changing density and overlap of the individual data points, estimation of the main trend would be greatly complicated and plausible error bars would be effectively impossible to calculate.

Our general statistical framework in this analysis was to assume that some unknown “canonical function” describes the average change in the feature-scaled GABA signal over the lifespan as a function of age. In other words, given participant *i* in study *s*, their age is denoted by *x_s,i_* and the relative change in GABA over the lifespan is given by the function *g* (*x_s,i_*). This change cannot be directly observed though imaging, but is instead inferred indirectly from a measurable reference (in our case, either water or creatine). As such, the mean observed effect in each study has an unknown feature scaling factor *F_s_*. Finally, individual observations are assumed to vary with respect to the lifespan function given normally-distributed noise *∊* with an unknown error term *σ*. In total, this gives the following general form for each observation in our data *y_s,i_*:

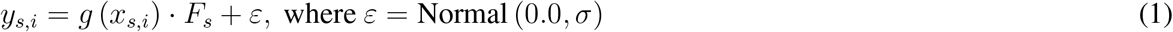

Because the scaling factor *F_s_* is intended to act as a feature-scaled standardization of the unitless function *g* (*x_s,i_*), the scaling factors were constrained so that the geometric mean of all values *g* (*x_s,i_*) were equal to 1.0. In addition to allowing different reference methods to be combined, the relative values of *F_s_* act as a study-by-study correction for any systematics that might otherwise shift one study’s observations relative to another’s.

This model is highly general, accommodating any function *g*() the analyst deems appropriate. The parameters that must be estimated are one scaling factor *F_s_* for each study, a global error term *σ*, and whichever parameters the function *g*() requires to specify its shape. Because parameter estimates in any non-linear regression model necessarily covary, it is essential that all parameters be estimated simultaneously (McElreath, 2020). If, for example, each dataset were “feature scaled” independently and then subsequently stitched together, any vertical shift needed to maximize the overlap of outcomes recorded during overlapping age ranged would necessarily be ad-hoc and would not be able to balance the relative weight of the evidence from each study in that area of overlapping age. By this same token, it was important that all studies included in our analysis included a range of ages that overlapped with at least one other study.

In order to ensure that all estimates were permitted to covary appropriately, we obtained posterior distributions for each parameter numerically using a Bayesian paradigm (Gelman et al., 2014) and implemented with the Stan programming language (Carpenter et al., 2017). Details of these analyses can be found in the supplementary information.

## RESULTS

To evaluate the 8 datasets for a linear trend, each dataset was the subject of a separate linear regression with respect to age. In each case, data were scaled by dividing the values by that dataset’s geometric mean. The resulting fits are depicted in Figure 3, with corresponding regression statistics reported in Table 3. These results are not meta-analytic; instead, they reflect the linear trend in each dataset. In six of the eight datasets, the linear trend explained less than 20% of the variance, and although the slope in the dataset with the youngest participants was positive, slopes tended to become more negative as the age of the participants increased. This pattern suggests that although a linear trend may provide a good account of data over short periods of time, a linear trend over the entire lifespan is not appropriate.

To provide a meta-analytic synthesis of these datasets without biasing our result through an arbitrary choice of our function *g*(), we fit two models: both nonlinear and able to accommodate the pattern visible from the individual trendlines.

Our first function *g* () was a penalized basis spline model, adapting the procedure described by Kharratzadeh (2017). This provided a nonlinear and nonparametric estimate of how GABA changes over the lifespan as described by these 8 datasets. The resulting model is depicted in Figure 4 (left). Overall, this time course is characterized by a rapid increase in GABA during early life, followed by a plateau from adolescence through midlife, and then a gradual decline from approximately 40 years onward.

**Figure 4.**
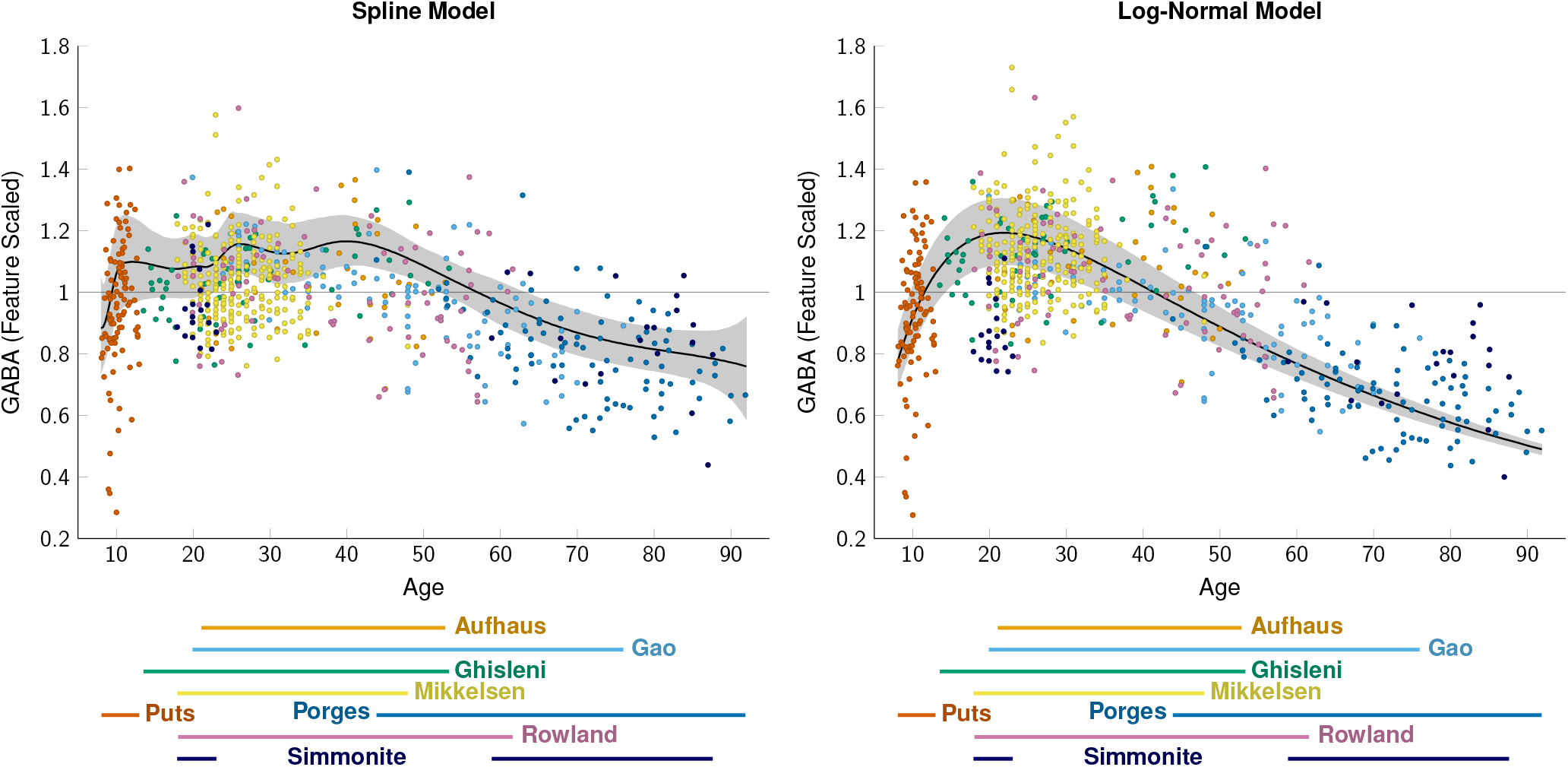
Nonlinear regression models of GABA signal integrating all data simultaneously. The shaded region depicts the 95% credible interval for the mean. **Left.** Penalized B-spline model. **Right.** Log-normal model.

This pattern of a rapid increase in GABA in early life, followed by a gradual decline, has been characterized in the past using the log-normal distribution to describe other neurophysiological changes across the lifespan (Lebel et al., 2012). To provide a parametric description of GABA over the lifespan, our second function g() was a log-normal distribution with a free scaling factor. This model is depicted in Figure 4 (right).

To ensure that the directionality of the ‘late-age’ component was not driven by changes in creatine, we performed an additional ‘leave-one-out’ (LOO) cross-validation approach (Arlot and Celisse, 2010) for each of the three datasets that included late-age participants. Figure 5 shows that when datasets are left out, the general shape of both models remains the same, with wider 95% credible intervals for the oldest of age due to limited available data. This suggests the overall pattern of a negative slope in late age is unlikely to be driven by creatine changes, as the data contributed by Porges et al. (2017b) made the largest contribution to this slope (having the most observations to contribute) and relied on a water-based reference method. We also explored the potential effect of region by performing the analysis on only frontal data (removing Mikkelsen et al. and Simmonite et al. respectively) with no substantial impact on the direction of the slope over time. This analysis is depicted in Figure 6.

**Figure 5.**
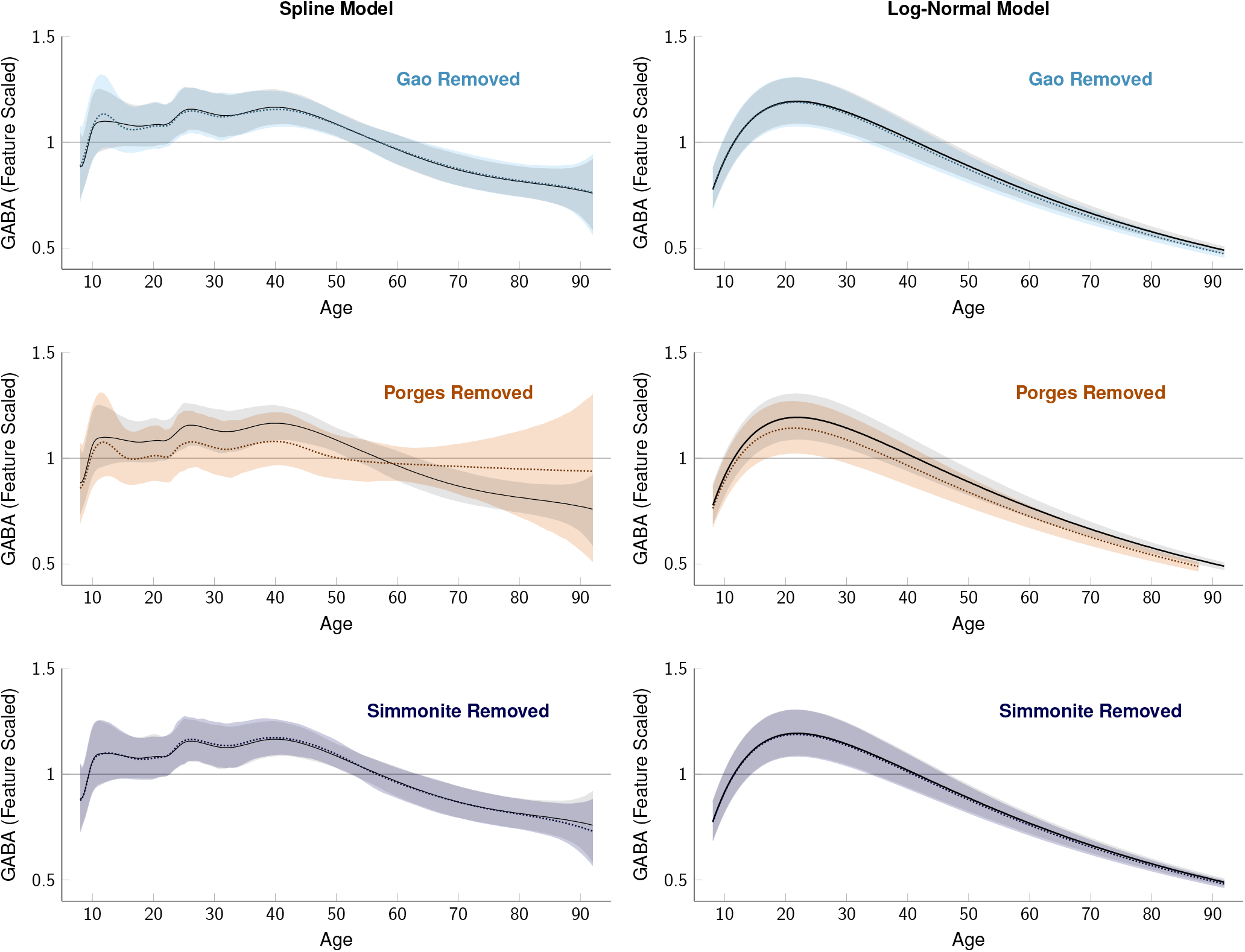
Nonlinear regression models of GABA signal performed using a leave-one-out (LOO) cross-validation approach for late-life datasets. The grey shaded region and black line depicts the 95% credible interval for the mean for the model as shown in Figure 4 and the colored shaded region and dotted line show the model with the respective data left out. In all cases, a similar overall trajectory to the full data is implied by each of the subsets, albeit with greater uncertainty.

**Figure 6.**
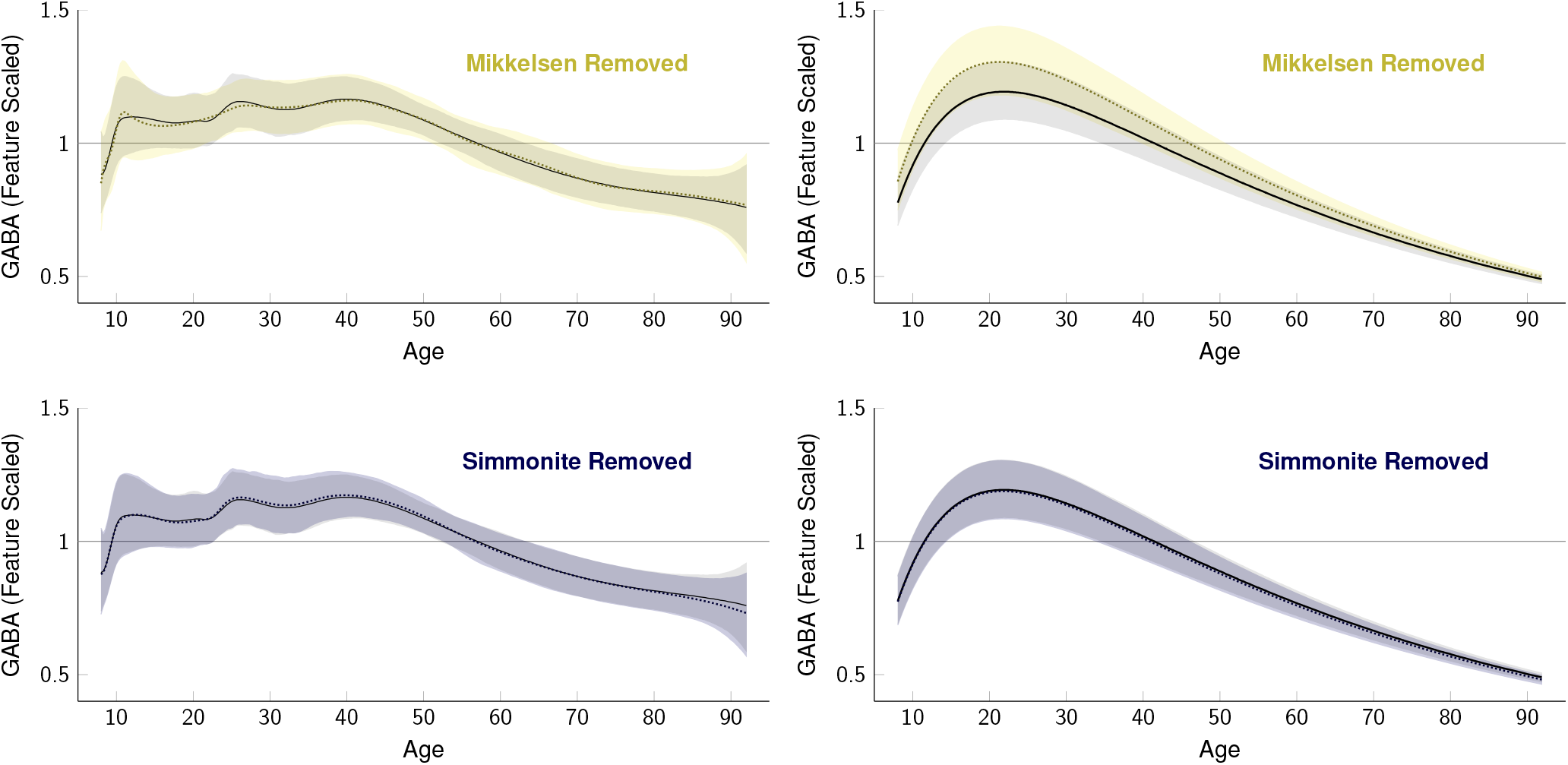
Nonlinear regression models of GABA signal performed using a leave-one-out (LOO) cross-validation approach for non-frontal datasets. The grey shaded region and black line depicts the 95% credible interval for the mean for the model as shown in Figure 4 and the colored shaded region and dotted line show the model with the respective data left out. In all cases, a similar overall trajectory to the full data is implied by each of the subsets, albeit with greater uncertainty.

## DISCUSSION

Here we show that the age-related trajectory of GABA across the lifespan is characterized by a sharp increase in GABA levels during development, followed by a flattening during early adulthood and by a subsequent slow decrease with aging. No single study reveals such a relationship (as evidenced by Figure 3) — it is only by meta-analysis of multiple datasets that we are able to identify this relationship. Here, we will discuss the case for a biological mechanism that drives a non-linear trajectory of GABA levels with age and then discuss methodological and biological factors that may influence these results. Finally, we review the implications of our evaluation and suggest potential directions of future research characterizing the trajectories of GABA concentrations over the lifetime.

### Biological Mechanisms

Although previous work, including the authors’ own, has generally reported a linear relationship between GABA and age within a given stage of life, it is far more plausible that changes across the lifespan are non-linear, in line with other biological effects. For example, while Gao et al. (2013) might capture a decrease in GABA with age, the broad age range studied in Ghisleni et al. (2015) might prevent observation of a linear relationship. Indeed, a decline in GABAergic interneurons with age has been widely reported in animal models (Hua et al., 2008; Stanley et al., 2012). Post-mortem data from human samples showed a reduction in the 65 isoform of Glutamic Acid Decarboxylase (the enzyme responsible for the production of GABA, GAD) in visual cortex, suggesting reduced GABA production with aging (Pinto et al., 2010). MRS of GABA by itself cannot provide sufficient resolution to determine what these reductions in GABA levels reflect; we can only make vague assumptions and conclusions on the relationship between brain structure, cognitive function, and potential molecular mechanisms.

In considering a nonlinear model for GABA across the human lifespan, we find three major stages: 1) a developmental stage where GABA levels increase rapidly, 2) a stabilization phase during adulthood where GABA concentrations remain mostly stable, and 3) a gradual descending period of GABA with advanced aging. This is consistent with previous studies of brain structural and cognitive function across the lifespan. Non-linear trajectories have been reported in age-effects of total grey matter (Lenroot et al., 2007; Sussman et al., 2016) and cortical thickness (Shaw et al., 2008). Diffusion tensor imaging (DTI) studies show non-linear age-related changes through childhood and adolescence (Lebel and Beaulieu, 2011), with an “inverted U-shaped” trajectory that peaks at approximately 40 years of age (Bendlin et al., 2010; Good et al., 2001; Lebel et al., 2012; Westlye et al., 2010). A similar trajectory pattern for cognitive abilities is reported in memory, verbal ability, and inductive reasoning (Kobayashi et al., 2015), as well as word recall, verbal fluency, math skills (Whitley et al., 2016), and behavioral inhibition (Williams et al., 1999).

#### GABA changes in Development

An increase in cortical GABA could be extrapolated from the proliferation of new GABAergic neurons during development. GABA is thought to be linked to the myelination of frontal white matter trajectories by controlling oligodendrocyte precursor cell activity through the developmental phase (Ghisleni et al., 2015; Vélez-Fort et al., 2012). An increase in cortical GABA concentrations could also be the result of increased synaptic activity of GABAergic neurons during development. This potentiation-like mechanism of cortical GABA is supported by studies where astrocytes have been shown to mediate plasticity of rodent hippocampus (Kang et al., 1998) and visual cortex (Chen et al., 2012), suggesting a possible potentiation-like mechanism of cortical GABA concentrations. In addition, upregulation of GAD could lead to increased production of GABA and indeed, GABA levels as measured with MRS have been shown to relate to expression of the GAD1 gene (Marenco et al., 2010). GABAergic neuronal function is reported to become more efficient through synaptic pruning and long-term depression during development (Paolicelli et al., 2011; Wagner and Alger, 1995; Wu et al., 2012). It should be pointed out that no eligible MRS studies of GABA were available for infancy and early development, which is a significant gap in the literature that should be addressed in future work.

#### GABA changes in Aging

Decreased GABA concentrations during aging are most likely linked to grey matter atrophy and demyelination associated with pathology and normal aging. Studies have shown a reduction in the number of interneurons expressing GAD in the medial prefrontal cortex. These reductions were accompanied by altered spatial working memory, linking altered GABA function to altered behavior in aging (Spiegel et al., 2013) (Spiegel et al., 2013). Furthermore, studies have shown changes in the efficacy of GABAergic function at both GABA-A and GABA-B receptors (McQuail et al., 2015). Finally, animal work has shown reduced GABAergic neurons and efficacy in cats and monkeys (Hua et al., 2008; Leventhal et al., 2003). Any other explanation of decreased cortical GABA during development may be linked to disease pathologies.

#### GABA changes across the lifespan

Very few non-MRS studies have assessed changes in GABA function across the lifespan. In those that report changes in the GABAergic system across the lifespan, a pattern of change similar to our findings has been presented. Using GAD labeling methodology in the human visual cortex, GAD65 has been reported to increase early in life and gradually decrease during aging (Pinto et al., 2010). Provocatively, they show GAD67 to be stable across the lifespan. Given the more specific relationship between GAD65 and neurotransmission, this may underlie reports of GABA associated alterations in cognitive function that occur during periods of GABAergic change (Porges et al., 2017b) throughout the lifespan. Future studies should aim to address the relationship between these two different functional isoforms, brain GABA levels, and function in both health and disease.

#### Regional differences

It can be assumed that regional changes in GABA are not homogeneous. We do not presume that the model we have presented is characteristic of age-related changes in GABA in all neural tissues due to well-known ontological and age-related regional variation in tissue that influence GABA levels (Lebel et al., 2012). One could even argue that regional differences are likely to exist within smaller regions of the frontal lobe. However, limiting this regional selectivity even further would have not allowed for our analysis, as 1) we would not be able to correct for covariation between frontal and parietal regions, and 2) Inclusion of multiple observations per participant would violate the independence of individual datapoints in our analysis. Because all data we included from a particular study were restricted to the same cortical region, differences between regions would be absorbed by the studies’ scaling factors *F_s_*, and would be indistinguishable from methodological systematics (further discussed below). We additionally show that the model shows a similar trajectory for frontal-only data (Figure 6). The majority of published studies on MRS of GABA focus on cortical, rather than subcortical, regions of interest. The few publications that describe subcortical GABA often show trends that are difficult to associate with cortical GABA levels. For instance, GABA was negatively associated with age in subcortical voxels but positively associated with age in anterior and posterior cortical voxels (Ghisleni et al., 2015). Subcortical GABA undoubtedly plays an important role in brain function, as evidenced by increased GABA concentrations in subcortical basal ganglia that have been associated with schizophrenia and depression Puts and Edden (2012). However, the role of subcortical GABA—measured via MRS—in cognitive function is not thoroughly investigated in the literature, and thus there are too few studies to perform a suitable meta-analysis. In the current review, we limit our discussion to GABA concentrations in cortical voxels in order to provide a meaningful initial evaluation of GABA over the lifespan. Both the publication of future participant-level data and the release of participant-level data from past studies will increase the sample available to the field, which in turn will make possible the rigorous examination of the contributions of these covariates.

### Methodological Differences

Inconsistencies in age-related changes in GABA levels may stem from differences between study methodologies or inherent structural differences across the lifespan. We discuss each of these in turn below.

Quantification of MRS is relative and expresses the ratio between the signal of interest and an internal reference signal. The most widely used references for GABA levels are the Cr signal in editing-off spectra and the unsuppressed water signal from the same volume (Alger, 2010; Mullins et al., 2014). Each method has its advantages and disadvantages. For instance, Cr is acquired during the same MEGA-PRESS scan as GABA, limiting effects of chemical shift: Cr has a minimal shift from the GABA signal. In contrast, the water signal represents a more concentrated chemical yielding a higher SNR, but it may also introduce error in estimates of location due to chemical shift effects (Mullins et al., 2014). Furthermore, the Cr signal arises only from tissue, whereas water signal arises from tissue and CSF with substantially different relaxation behavior. A small number of studies have looked at the relationship between Cr and age (Ding et al., 2016) showing an increase or no change with age (for review, see Cleeland et al., 2019). This would make the negative GABA/Cr correlation less significant. Moreover, Gao et al. investigated both GABA/Cr and GABA/NAA and found the same effect. NAA has been shown to consistently decrease in aging as well (Cleeland et al., 2019). There remains the potential for a Cr-bias on the final model that needs to be considered. Our leave-one-out approach to the ‘late-age’ data (Figure 5) found no substantial differences in the final models, providing additional support to the notion that differences in creatine are not driving the GABA findings presented here.

Another limitation is that voxel location can be inconsistent between studies (e.g. the medial prefrontal cortex in one study may not be localized the same way as in another study, (also see a recent review Peek et al., 2020). This becomes more problematic in younger cohorts where, due to smaller intracranial volumes, the methodologically limited size of the voxel (Mullins et al., 2014) necessarily incorporates a proportionally larger fraction of the brain.

It is well known that voxel tissue composition has a significant impact on the quantification of GABA levels (Harris et al., 2015). In many cases, tissue correction is appropriate due to existing partial volume effects (Barker et al., 1993; Christiansen et al., 1993; Danielsen and Henriksen, 1994; Ernst et al., 1993; Hennig et al., 1992; Kreis et al., 1993; Thulborn and Ackerman, 1983). Researchers frequently tissue-correct GABA values by segmenting the T1 weighted structural images (Ghisleni et al., 2015). Interestingly, none of the studies included in this review showed significant changes in segmented tissue content with age. This is surprising because other studies (Maes et al., 2018; Porges et al., 2017a) showed that age-related changes in GABA depend on tissue correction due to atrophy being common in older participants. The potential effect of tissue composition on GABA levels is limited if a study focused on a young cohort where age-related atrophy would be negligible. All of the studies included in this review assume a linear model to represent change in GABA. Thus, while tissue correction could potentially contribute to the variance and differences between MRS-GABA studies, they are unlikely to explain the discrepancies between study populations of different age ranges.

In water-referenced studies of age-related changes in older cohorts (where bulk tissue changes are most likely to impact this relationship), accounting for cerebrospinal fluid (CSF) in the analysis does not remove a significant relationship between age and cortical GABA (Porges et al., 2 017a). Given our concern was with relative changes in GABA, our analysis approach used a feature-scaling approach to examine variation within-study as an estimation problem that was simultaneous with the estimation of the time course. Given this approach, the scaling factor *F_s_* for each study s is conceptually very similar to a method of correcting “house effects” in the analysis of political polling data (Jackman, 2005), allowing pollsters that show systematic bias (e.g. to a particular party) to be included in aggregate measures of public opinion. Indeed, there is no significant difference in the correlation for GABA x age in the overlapping age range of 43 to 76 years between Porges et al. (GABA+/H2O) and Gao et al. (GABA/Cr) who looked at comparable samples (Fisher R-to-Z; (z = 1.48, p = 0.19)) showing consistency between studies focusing on the same population.

As for other methodological differences (Table 1 and Table 2), studies used a variety of scanner vendors: a variation that is known to contribute to between-site variability (Mikkelsen et al., 2017) but is unlikely to lead to substantial difference in the age relationship. Furthermore, a variety of analysis techniques have been used, making direct comparison between studies problematic. A given analysis pipeline is unlikely to be biased to substantial differences in the age relationship. By rescaling the data in a way that emphasized scale ratios within each dataset, we minimized the impact of differences in site and vendor-driven GABA magnitude estimations. Future studies should provide concentration values across the lifespan. Recent developments in MRS of GABA methodology will allow for this, even when data is collected on multiple scanner platforms (Saleh et al., 2020). It should be noted that all datasets used for this meta-analysis were cross-sectional. Although it is tempting to draw longitudinal inferences from our analysis, there are several limitations in doing so. For example, the present data provide no insight into long-term survival trends, so the population that is represented at age 20 likely differs in various ways from the population that is represented at age 70 and we cannot rule out a relationship between GABA and survival-related confounds. Additionally, because our method simultaneously estimated the scaling factor and the time course for each dataset, areas of minimal overlap between datasets (e.g. at around 13 years) are particularly uncertain.

The present analysis provides a model of the lifespan based on the data that is presently available. It would be valuable to test these in a longitudinal study of MRS of GABA across the lifespan. Even if it is not feasible to measure GABA in the same individual over a 70-year span, obtaining multiple estimates per participant over a moderate length of time (e.g. 5 years) would greatly facilitate estimating rate of change over time.

## Conclusion and Future Research

This review considered research of cortical GABA concentrations over the lifespan. After combining datasets, we conclude that a linear model of GABA over the lifetime is not supported. Instead, consistent with other developmental aging studies of neurophysiology and cognitive function, we propose non-linear models to describe lifespan GABA levels. A log-normal trajectory provides a satisfactory parametric description of the life-course for the time being, but large and longitudinal datasets may necessitate the use of nonparametric regression strategies to best characterize the age-GABA relationship.

In the future, it will be important to investigate the neurophysiological and anatomical processes that drive apparent changes in bulk metabolite and neurotransmitter levels. While it is clear that GABA changes with age and seems to follow trends reported in other lifespan datasets, non-linear relationships in GABA and other neurotransmitter concentrations warrant further exploration. Future inquiries would benefit from recruiting cohorts that encompass the entire lifespan. Studies of development including those of infants and young children are virtually non-existent but are crucial given the importance of GABA in early development. Furthermore, alterations in GABA have been seen in neurodevelopmental and neurodegenerative disorders—a better understanding of abnormal GABAergic function early in development may elucidate this relationship, point to potential early-intervention targets, or explain variability in response to pharmacological treatments for these conditions.

Here, we report on the relationship between GABA and age across the lifespan. It is well known that GABA contributes to cognition and perception, which in and of themselves change with age. It would be extremely interesting to apply our meta-analytical approach to the investigation of how age-related changes in GABA might correlate with cognition. However, this is greatly complicated by the wide variety of cognitive measures used across studies, which would be much more difficult to standardize across studies than feature-scaled GABA as measured using an MRI machine. While pursuit of this work is notably challenging, conducting such research is undeniably crucial going forward.

While many papers exist that report on age-related changes that are consistent with our lifespan trajectory, these are often reported as group or cohort differences without the presentation of individual data points necessary for such meta-analyses as applied here (Hermans et al., 2018; Port et al., 2017). Collaborative and group-science approaches are becoming increasingly important in generating large datasets that allow for a broader and larger-scale application of this work and we hope that sharing data, or at least reporting individual data points, becomes more common in the future even in studies where age may not be the main focus.

## Supporting information

Data & Analysis

