## Supplementary material for "The trajectory of cortical GABA levels across the lifespan: An individual participant data meta-analysis of edited MRS studies": Data & Analysis: GABASpline.R

```
library(rstan)
library(splines)
library(rethinking)
library(readr)

# Change source_folder to appropropriate target folder
source_folder <- "~" 

#================================
#==== MRS Gaba Data Analysis ====
#================================

#====Assembling Data====

data <- list()

data[1] <- list(read_csv(paste(c(source_folder,"Data/Auf_f_fs.csv"),collapse = ''), col_names=FALSE))
data[2] <- list(read_csv(paste(c(source_folder,"Data/Gao_f_fs.csv"),collapse = ''), col_names=FALSE))
data[3] <- list(read_csv(paste(c(source_folder,"Data/Ghis_f_fs.csv"),collapse = ''), col_names=FALSE))
data[4] <- list(read_csv(paste(c(source_folder,"Data/Mik_f_fs.csv"),collapse = ''), col_names=FALSE))
data[5] <- list(read_csv(paste(c(source_folder,"Data/Por_f_fs.csv"),collapse = ''), col_names=FALSE))
data[6] <- list(read_csv(paste(c(source_folder,"Data/Puts_fr_fs.csv"),collapse = ''), col_names=FALSE))
data[7] <- list(read_csv(paste(c(source_folder,"Data/Row_f_fs.csv"),collapse = ''), col_names=FALSE))
data[8] <- list(read_csv(paste(c(source_folder,"Data/Simm_p_fs.csv"),collapse = ''), col_names=FALSE))

X <- c()
Y <- c()
cat <- c() 
C <- 8
for (i in 1:C) {
	X <- c(X,data[[i]]$X1)
	Y <- c(Y,data[[i]]$X2)
	cat <- c(cat,rep(i,length(data[[i]]$X1)))
}
N <- length(X)
plot(X,Y)

cyc <- 2000

rstan_options(auto_write = TRUE);
options(mc.cores = parallel::detectCores());

cols <- c("red", "chocolate", "darkgoldenrod1", "blue", "purple", "deeppink", "chartreuse3", "navyblue")

#====Naive Linear Fits====

simple_linear_fits <- list()
simple_linear_model<-stan_model(paste(c(source_folder,"Stan/linear_simple.stan"),collapse = ''))
mns <- matrix(0,C,2)
outX <- seq(8,99,by=0.1)
outY <- matrix(0,length(outX),C)
for (i in 1:C) {
	dex <- (cat==i)
	x <- X[dex]
	y <- Y[dex]
	mns[i,1] <- mean(x)
	mns[i,2] <- mean(y)
	dat <- list(N=sum(dex), X=x-mns[i,1], Y=y/mns[i,2])
	simple_linear_fits[i] <- list(sampling(simple_linear_model, data=dat, iter=cyc*2, warmup=cyc, chains=4, cores=4, control = list(adapt_delta=0.999,max_treedepth=15)))
}
plot(c(NA),c(NA), typ="l", lwd=3, xlim=c(8,100), ylim=c(0,3),xlab="Age",ylab="GABA (Feature Scaled)",col=cols[i])
for (i in 1:C) {
	dex <- (cat==i)
	x <- X[dex]
	y <- Y[dex]
	mns[i,1] <- mean(x)
	mns[i,2] <- mean(y)
	q <- extract.samples(simple_linear_fits[[i]])
	Y_fit <- matrix(0,cyc*4,length(outX))
	for (j in 1:(cyc*4)) {
		Y_fit[j,] <- (q$beta[j,1] + q$beta[j,2]*(outX-mns[i,1]))
	}
	yf <- apply(Y_fit,2,mean)
	yf.PI <- apply(Y_fit,2,PI,prob=0.95)
	lines(outX,yf,col=cols[i])
	shade(yf.PI, outX, col=alpha(cols[i],0.4))
	points(x,y/mns[i,2], cex=0.5, pch=19,col=cols[i])
}

v <- rep(0,8)
for (i in 1:C) {
	dex <- (cat==i)
	x <- X[dex]
	y <- Y[dex]
	mns[i,1] <- mean(x)
	mns[i,2] <- mean(y)
	q <- extract.samples(simple_linear_fits[[i]])
	Y_fit <- matrix(0,cyc*4,length(x))
	for (j in 1:(cyc*4)) {
		Y_fit[j,] <- (q$beta[j,1] + q$beta[j,2]*(x-mns[i,1]))
	}
	yf <- apply(Y_fit,2,mean)
	v[i] <- var(yf)/var(y/mns[i,2])
}


#====Preparing & Fitting The Spline====

spline_factor_penalized_model <- stan_model(paste(c(source_folder,"Stan/b_spline_factor_penalized.stan"),collapse = ''))
K <- 19                                                               # Number of knots
D <- 3                                                                # Spline degree
num_basis <- K + D - 1                                                # Number of basis splines
knots <- unname(quantile(X,probs=seq(from=0, to=1, length.out = K)))  # Uniform knot placement
dat <- list(N=N, K=K, C=C, knots=knots, D=D, cat=cat, X=X, Y=Y)
fit_spline <- sampling(spline_factor_penalized_model, data=dat, iter=cyc*2, warmup=cyc, chains=4, cores=4, control = list(adapt_delta=0.999,max_treedepth=15))
precis(fit_spline,pars=c("a0","sigma","a_tau"),depth=2)

#====Interpreting The Model====

B <- t(bs(X, df=num_basis, degree=D, intercept = TRUE))               # Basis matrix
q <- extract.samples(fit_spline)

Y_fit <- q$a%*%B
crrt <- mean(Y_fit)
for (i in 1:(cyc*4)) {
	Y_fit[i,] <- Y_fit[i,]/mean(Y_fit[i,])
}
x <- sort.int(X,index.return = TRUE)
yf <- apply(Y_fit,2,mean)[x$ix]
yf.PI <- apply(Y_fit,2,PI,prob=0.95)
plot(x$x,yf, typ="l", lwd=3, ylim=c(0,2),xlab="Age",ylab="GABA (Feature Scaled)",main="Spline Fit")
shade(yf.PI[,x$ix], x$x)

for (i in 1:C) {
	points(X[cat==i],(Y[cat==i]/mean(q$a0[,i]))/crrt, cex=0.5, pch=19,col=cols[i])
	subx <- X[cat==i]
	dex <- (x$x>=min(subx))&(x$x<=max(subx))
	
	yh <- apply(Y_fit[,x$ix],2,mean)
	lines(x$x[dex],yh[dex], lwd=1, col=cols[i])
}

lbls <- c("Auf","Gao","Ghis","Mik","Por","Puts","Row","Simm")
boxplot(q$a0/crrt,outline=FALSE,names=lbls,ylab="Multiplicative House Effect")

#====Lognormal Models====

lognormal_model<-stan_model(paste(c(source_folder,"Stan/lognormal_factor.stan"),collapse = ''))
fit_lognormal <- sampling(lognormal_model, data=dat, iter=cyc*2, warmup=cyc, chains=4, cores=4, control = list(adapt_delta=0.999,max_treedepth=15))
precis(fit_lognormal,pars=c("loc","disp","a0","sigma"),depth=2)
q <- extract.samples(fit_lognormal)

Y_fit <- matrix(0,cyc*4,N)
for (i in 1:(cyc*4)) {
	Y_fit[i,] <- dlnorm(X,q$loc[i],q$disp[i])
}
crrt <- mean(Y_fit)
Y_fit <- Y_fit/crrt
x <- sort.int(X,index.return = TRUE)
yf <- apply(Y_fit,2,mean)[x$ix]
yf.PI <- apply(Y_fit,2,PI,prob=0.95)
plot(x$x,yf, typ="l", lwd=3, ylim=c(0,2),xlab="Age",ylab="GABA (Feature Scaled)",main="Log-normal Fit")
shade(yf.PI[,x$ix], x$x)

for (i in 1:C) {
	points(X[cat==i],(Y[cat==i]/mean(q$a0[,i]))/crrt, cex=0.5, pch=19,col=cols[i])
	subx <- X[cat==i]
	dex <- (x$x>=min(subx))&(x$x<=max(subx))
	yh <- apply(Y_fit[,x$ix],2,mean)
	lines(x$x[dex],yh[dex], lwd=1, col=cols[i])
}
```
