## Supplementary material for "The trajectory of cortical GABA levels across the lifespan: An individual participant data meta-analysis of edited MRS studies": Data & Analysis: b_spline_factor_penalized.stan

```
functions {
	vector build_b_spline(real[] t, real[] ext_knots, int ind, int order);
	vector build_b_spline(real[] t, real[] ext_knots, int ind, int order) {
		// INPUTS:
		//    t:          the points at which the b_spline is calculated
		//    ext_knots:  the set of extended knots
		//    ind:        the index of the b_spline
		//    order:      the order of the b-spline
		vector[size(t)] b_spline;
		vector[size(t)] w1 = rep_vector(0, size(t));
		vector[size(t)] w2 = rep_vector(0, size(t));
		if (order==1) {
			for (i in 1:size(t)) { // B-splines of order 1 are piece-wise constant
				b_spline[i] = (ext_knots[ind] <= t[i]) && (t[i] < ext_knots[ind+1]);
			}
		} else {
			if (ext_knots[ind] != ext_knots[ind+order-1]) {
				w1 = (to_vector(t) - rep_vector(ext_knots[ind], size(t)))/(ext_knots[ind+order-1] - ext_knots[ind]);
			}
			if (ext_knots[ind+1] != ext_knots[ind+order]) {
				w2 = 1 - (to_vector(t) - rep_vector(ext_knots[ind+1], size(t)))/(ext_knots[ind+order] - ext_knots[ind+1]);
			}
			// Calculating the B-spline recursively as linear interpolation of two lower-order splines
	    	b_spline = w1 .* build_b_spline(t, ext_knots, ind, order-1) + w2 .* build_b_spline(t, ext_knots, ind+1, order-1);
		}
		return b_spline;
	}
}
data {
	int<lower=0> N;         // Number of data points
	int<lower=1> K;         // Number of knots
	int<lower=0> C;         // Number of data categories
	vector[K] knots;        // the sequence of knots
	int<lower=1> D;         // the degree of spline (is equal to order - 1)
	int<lower=0> cat[N];    // Data category index 
	real Y[N];              // Observed outcome values
	real X[N];              // Observed predictor values
}
transformed data {
	int num_basis = K + D - 1;           // Total number of B-splines
	matrix[num_basis, N] B;              // Matrix of B-splines
	vector[D + K] ext_knots_temp;
	vector[2*D + K] ext_knots;           // Set of extended knots

	ext_knots_temp = append_row(rep_vector(knots[1], D), knots);
	ext_knots = append_row(ext_knots_temp, rep_vector(knots[K], D));
	for (i in 1:num_basis) {
		B[i,:] = to_row_vector(build_b_spline(X, to_array_1d(ext_knots), i, D + 1));
	}
	B[K + D - 1, N] = 1;
}
parameters {
	real<lower=0> a0[C];			// House effect (per category)
	row_vector[num_basis] a_raw;	// Moving center of mass
	real<lower=0> a_tau;			// Smoothing factor for a
	real<lower=0> sigma;			// Uniform dispersion
}
transformed parameters {
	row_vector[num_basis] a;        
	vector[N] Y_hat;

	a[1] = a_raw[1];
	for (i in 2:num_basis) {
		a[i] = a[i-1] + a_raw[i]*a_tau;
	}
	Y_hat = to_vector(a*B);
	for (n in 1:N) {
		Y_hat[n] = Y_hat[n]*a0[cat[n]];
	}
}
model {
	// Priors
	a0 ~ normal(1, 2);
	a_raw ~ normal(0, 0.1);
	a_tau ~ exponential(0.1);
	sigma ~ exponential(1);
	//Likelihood
	Y ~ normal(Y_hat, sigma);
}
generated quantities {
	vector[N] log_lik;
	for ( n in 1:N ) {
		log_lik[n] = normal_lpdf(Y[n] | Y_hat[n], sigma);
	}
}
```
