## Supplementary material for "The trajectory of cortical GABA levels across the lifespan: An individual participant data meta-analysis of edited MRS studies": Data & Analysis: linear_simple.stan

```
data {
	int<lower=0> N;         // Number of data points
	real Y[N];              // Observed outcome values
	real X[N];              // Observed predictor values
}
parameters {
	real beta[2];			// Intercept & Slope
	real<lower=0> sigma;	// Uniform dispersion
}
model {
	vector[N] Y_hat;
	// Priors
	beta ~ normal(0, 2);
	sigma ~ exponential(1);
	for (n in 1:N) {
		Y_hat[n] = (beta[1] + beta[2]*X[n]);
	}
	//Likelihood
	Y ~ normal(Y_hat, sigma);
}
```
