## Supplementary material for "The trajectory of cortical GABA levels across the lifespan: An individual participant data meta-analysis of edited MRS studies": Data & Analysis: lognormal_factor.stan

```
data {
	int<lower=0> N;         // Number of data points
	int<lower=0> C;         // Number of data categories
	int<lower=0> cat[N];    // Data category index 
	real Y[N];              // Observed outcome values
	real X[N];              // Observed predictor values
}
parameters {
	real<lower=0> a0[C];			// House effect (per category)
	real loc;						// Location
	real<lower=0> disp;				// Dispersion
	real<lower=0> sigma;			// Uniform dispersion
}
model {
	vector[N] Y_hat;
	// Priors
	a0 ~ normal(100, 40);
	loc ~ normal(4, 1);
	disp ~ exponential(1);
	sigma ~ exponential(1);
	for (n in 1:N) {
		Y_hat[n] = a0[cat[n]]*exp(lognormal_lpdf(X[n]|loc,disp));
	}
	//Likelihood
	Y ~ normal(Y_hat, sigma);
}
generated quantities {
	vector[N] log_lik;
	for ( n in 1:N ) {
		log_lik[n] = normal_lpdf(Y[n] | a0[cat[n]]*exp(lognormal_lpdf(X[n]|loc,disp)) , sigma);
	}
}
```
